# IL-2 immunotherapy rescues irradiation-induced T cell exhaustion *in vivo*

**DOI:** 10.1101/2024.01.29.577732

**Authors:** Carmen S. M. Yong, Irma Telarovic, Lisa Gregor, Miro E. Raeber, Martin Pruschy, Onur Boyman

**Affiliations:** Laboratory for Applied Radiobiology, Department of Radiation Oncology, University Hospital Zurich, University of Zurich, Zurich, Switzerland; Department of Immunology, University Hospital Zurich, Zurich, Switzerland; Faculty of Medicine and Faculty of Science, University of Zurich, Zurich, Switzerland

**Author notes:** Corresponding authors: Martin Pruschy, Ph.D., Laboratory for Applied Radiobiology, Department of Radiation Oncology, University Hospital Zurich, Raemistrasse 100; University of Zurich, 8006 Zurich, Switzerland.; Onur Boyman, M.D., Department of Immunology, University Hospital Zurich, Schmelzbergstrasse 26, 8091 Zurich, Switzerland. M.P. and O.B. contributed equally to senior authorship.

**Keywords:** radiotherapy, immunotherapy, interleukin-2

## Abstract

Radiotherapy (RT) can stimulate anti-cancer T cell responses that target primary and distant tumors. In addition to antigen-mediated stimulation of effector T cells, signals from stimulatory cytokines, notably interleukin-2 (IL-2), are necessary for optimal T cell function and memory. However, timing and IL-2 receptor (IL-2R) bias of such signals are ill-defined. Using image-guided RT in a mouse colon cancer model, we observed that single high-dose (1 x 20 Gy) RT transiently upregulated IL-2Rα (CD25) on effector CD8^+^ T cells, facilitating the use of CD25-biased IL-2 immunotherapy. Timed administration of CD25-biased IL-2 treatment after RT favored the expansion of tumor-infiltrating CD8^+^ T cells over regulatory T cells and IL-2Rβ (CD122)^high^ CD8^+^ T cells, which resulted in comparable anti-tumor effects as with RT plus CD122-biased IL-2 immunotherapy. Moreover, intratumoral CD8^+^ T cells from animals receiving combined IL-2R-biased IL-2 and RT showed reduced signatures of T cell exhaustion. Finally, these combination treatments affected both primary irradiated and distant non-irradiated tumors, achieving durable responses. We demonstrate that timed and IL-2R subunit-biased IL-2 immunotherapy synergized with single high-dose RT to achieve potent anti-cancer immunity.

## Introduction

Radiotherapy (RT) is a frontline treatment for cancer, inducing both direct and indirect mechanisms of tumor cell killing^1–3^. One major consequence of RT is its ability to stimulate an anti-cancer immune response. This is mediated via the induction of immunogenic cell death and the release of tumor antigens, along with pro-inflammatory factors, collectively resulting in the priming of antigen-specific T cells^4^. While the therapeutic effect of focal irradiation is mainly confined to the target area, in rare cases it has been described to induce systemic immune responses, resulting in control of non-irradiated, distant tumor lesions. This so-called ‘abscopal effect’ has been shown in pre-clinical models to be highly dependent on the dosage, fractionation regimen, and immunogenicity of tumor^5,6^. However, a mechanistic understanding is still lacking to reliably translate a respective RT regimen to the clinic. Of utmost importance, such an abscopal effect relies on an efficient and functioning immune system, in particular CD8^+^ T cells^6,7^. Even though a systemic immune response after RT alone is rarely robust enough to induce changes in non-irradiated lesions, this can be enhanced when combined with immunomodulatory agents^8^.

Interleukin-2 (IL-2) is a cytokine capable of stimulating both CD8^+^ T cells and natural killer cells (NK) as well as regulatory T cells (Tregs). Whereas Tregs express the trimeric form of IL-2 receptor (IL-2R), composed of IL-2Rα (CD25), IL-2Rβ (CD122) and IL-2Rγ (CD132), antigen-primed (memory) CD8^+^ T cells and NK cells express the dimeric form composed of CD122 and CD132^9–11^. IL-2 as a monotherapy has demonstrated low to moderate anti-tumor efficacy in patients with metastatic melanoma and metastatic renal cell cancer^12^. However, as a consequence of its short half-life, high doses of IL-2 are required to achieve substantial clinical responses, which in turn can result in unwanted and severe side effects, such as vascular leak syndrome^13–15^. Furthermore, the high affinity of trimeric IL-2Rs on Tregs outcompetes the dimeric IL-2Rs on CD8^+^ T cells, leaving the latter bereft of IL-2. To overcome these issues, different IL-2 variants have been developed with sustained *in vivo* half-life and biased targeting of IL-2 to either dimeric or trimeric IL-2Rs, which results in preferential expansion of specific immune cell subsets^10,11,16^. One such approach uses binding of unmodified IL-2 to particular anti-IL-2 monoclonal antibodies (mAb), which results in IL-2 complexes (IL-2cx)^16^. Such IL-2cx are either CD25-biased, stimulating CD25^high^ immune cells with high expression of trimeric IL-2Rs, or CD122-biased, activating CD122^high^ immune cells^15,17–23^. Use of CD122-biased IL-2cx resulted in potent anti-tumor immune responses in several mouse cancer models^20,23,24^.

The combination of RT with immunotherapy using high-dose unmodified IL-2 has shown promising results in preclinical and phase I and II clinical trials^25–29^. Moreover, RT was combined with IL-2 variants that preferentially targeted effector immune cells or antigens present in the tumor microenvironment^30–33^. Although these studies demonstrated some efficacy of combined RT and IL-2 immunotherapy, it is unknown which IL-2R signals and bias combine best with which RT treatment scheme. Notably, differences in dosage, fractionation regimens and conformity define the therapeutic efficacy of RT through direct damage of the irradiated tumor, but they also have a crucial impact on the anti-tumor immune response, particularly when combined with immunotherapy^34,35^.

Here, we studied the effects of hypofractionated RT and single high-dose RT on tumor-infiltrating T cells. We observed that single high-dose RT upregulated CD25 on intratumoral CD8^+^ T cells, which facilitated the combination of RT with a CD25-biased IL-2cx, resulting in efficient reduction of both primary irradiated and abscopal tumor nodules. Notably, RT alone induced a signature of exhaustion in tumor-infiltrating CD8^+^ T cells, which was reversed by use of IL-2cx immunotherapy, resulting in long-lived anti-tumor immune cell responses.

## Results

### Radiotherapy transiently upregulates CD25 on tumor-infiltrating CD8^+^ T cells

We first tested two previously published, isoeffective RT schemes, comparing locoregional irradiation with single high-dose RT of 20 Gy (1 x 20 Gy) versus a hypofractionated RT regimen of 8 Gy delivered over three consecutive days (3 x 8 Gy)^34^. C57BL/6 mice were injected subcutaneously with MC38 colon carcinoma cells, followed by tumor irradiation 10-11 days later, when tumors reached a size of ∼200 mm^3^ (**Fig. 1A**). We observed a distinct advantage in tumor control and long-term survival in mice treated with 1 x 20 Gy compared to mice receiving 3 x 8 Gy, as exemplified by 36% versus 8% complete remission, respectively (**Fig. 1B and 1C**).

**Figure 1.**
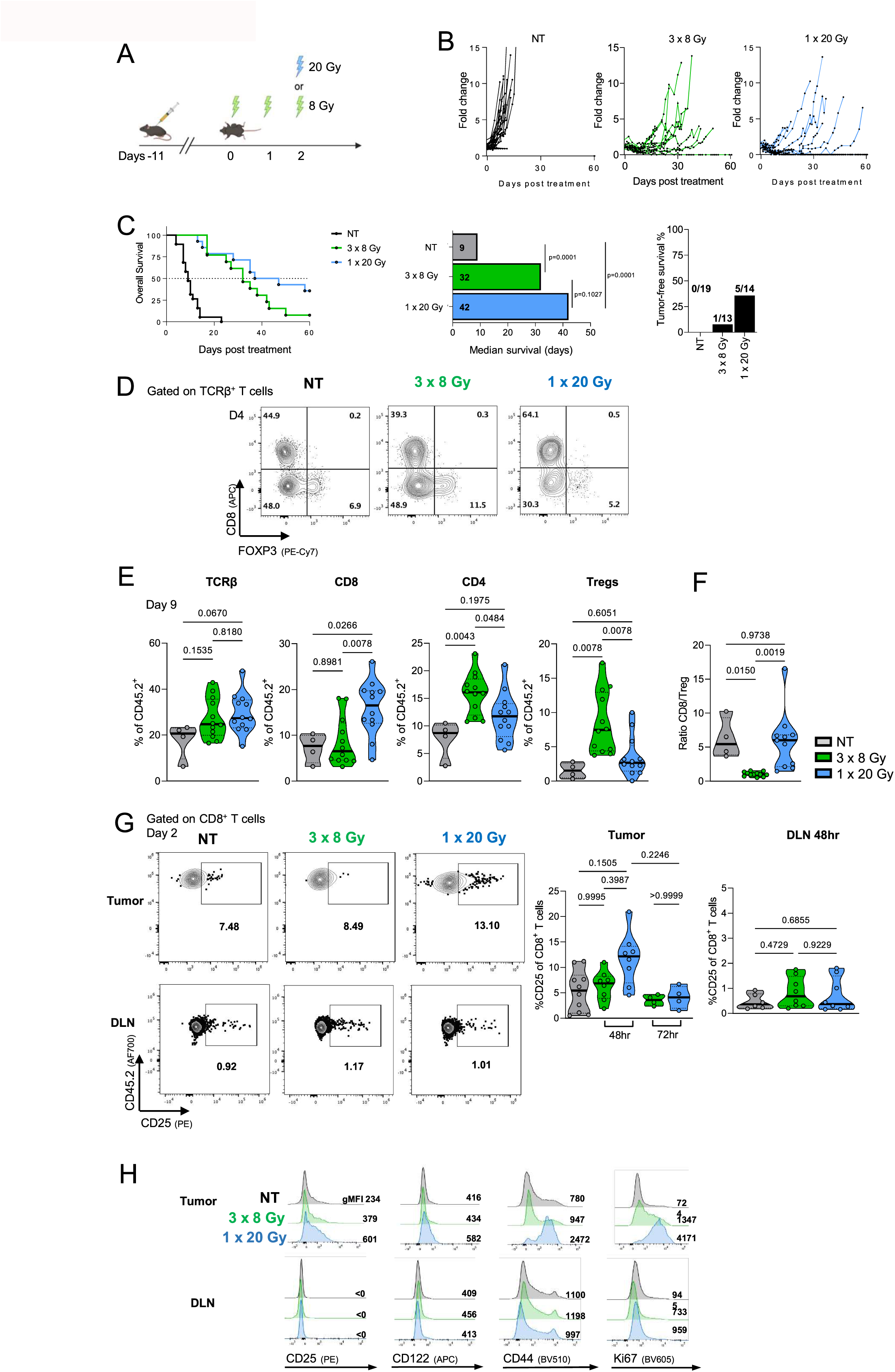
Single high-dose radiotherapy transiently upregulates CD25 on tumor-infiltrating CD8^+^ T cells. **(A)** Schematic representation of the experimental setup. C57BL/6 mice were injected subcutaneously with MC38 cells and tumors irradiated 10–11 days later. **(B)** Mice were treated with 8 Gy on three consecutive days or a single dose of 20 Gy. Tumor growth was monitored until day 60 post treatment. **(C)** Survival curve, median survival and percentage of tumor-free mice at experiment termination. **(D)** Representative flow cytometry plots gated on TCRβ^+^ T cells in the tumor and DLN four days after irradiation. **(E)** Percentages of TCRb^+^ T cells, CD8^+^ T cells, CD4^+^ T cells, and forkhead box P3 (FoxP3)^+^ regulatory T cells (Tregs) in the tumor 9 days after irradiation, and **(F)** ratios of CD8^+^ T cells to Tregs. **(G)** Representative flow cytometry plots gated on CD8^+^ T cells in tumor and tumor-draining lymph nodes (DLNs), (right) percentage of CD25^+^ cells in CD8^+^ T cells two days after irradiation. **(H)** Representative histograms of CD25, CD122, CD44 and Ki67 gated on CD8^+^ T cells in tumor and DLN four days after irradiation. Data are presented as mean ± standard error of the mean (SEM) of one to five independent experiments. **C)** Differences in median survival days were taken from the survival curves and analyzed by pairwise Log-rank (Mantel-Cox) test. **E-G)** Differences were analyzed using a one-way ANOVA.

By gating on T cell receptor β-expressing (TCRβ^+^) intratumoral T cell subsets, we determined that the proportion of CD8^+^ T cells and forkhead box p3 (Foxp3)^+^ Tregs was shifted in favor of CD8^+^ T cells in mice treated with 1 x 20 Gy (**Fig. 1D and 1E**). Conversely, mice treated with 3 x 8 Gy showed increased percentages of Foxp3^+^ cells and decreased percentages of CD8^+^ T cells compared to non-treated and 1 x 20 Gy-irradiated mice (**Fig. 1D and 1E**). Overall, these changes resulted in smaller CD8^+^ T cell-to-Treg (CD8/Treg) ratios in animals receiving 3 x 8 Gy compared to a single dose of 20 Gy (**Fig. 1F**).

We assessed phenotypic and functional changes, including IL-2R expression, in tumor-infiltrating CD8^+^ T cells following RT. On day 4 after RT, we observed a distinct upregulation of CD25, CD44 and the proliferation marker Ki67 on tumor-infiltrating CD8^+^ T cells preferentially in mice receiving 1 x 20 Gy, which contrasted with 3 x 8 Gy-treated and non-treated animals (**Fig. 1G and 1H**). Notably, the increase in CD25 was only observed in intratumoral CD8^+^ T cells, but not in Tregs (**Suppl. Fig. 1**, whereas their counterparts in tumor-draining lymph nodes (DLN) maintained low CD25 levels (**Fig. 1G and 1H**). Upregulation of CD25 occurred transiently after irradiation, with no differences in percentage between the treatment groups detectable on day 3 after RT. Although a slight increase in CD122 expression was also observed in tumor-infiltrating CD8^+^ T cells, but not in DLN CD8^+^ T cells, this effect was less apparent than the upregulation of CD25.

### Combined radiotherapy and IL-2 immunotherapy efficiently controls primary tumors and generates long-lasting anti-tumor immune responses

Having observed an upregulation of CD25 and, to a lesser degree, CD122 on tumor-infiltrating CD8^+^ T cells following a single dose of 20 Gy, we decided to combine 1 x 20 Gy with CD25-biased and CD122-biased IL-2cx, made of recombinant IL-2 complexed to the anti-IL-2 mAbs UFKA-20 and NARA1, respectively (**Fig. 2A**)^18,20^. Compared to non-treated animals, administering CD25-biased IL-2/UFKAcx (referred to as CD25-biased IL-2cx or IL-2cx_CD25_ from here on) alone only minimally affected tumor growth (**Fig. 2B**). However, combination of RT with either IL-2cx_CD25_ or CD122-biased IL-2/NARA1cx (referred to as CD122-biased IL-2cx or IL-2cx_CD122_ from here on) strongly delayed tumor growth (**Fig. 2B**). Combined RT plus IL-2cx treatment provided a significant survival advantage compared to RT alone (**Fig. 2C**). Furthermore, combined single high-dose RT plus IL-2cx immunotherapy resulted in 50% or more complete remission (**Fig. 2D**).

**Figure 2.**
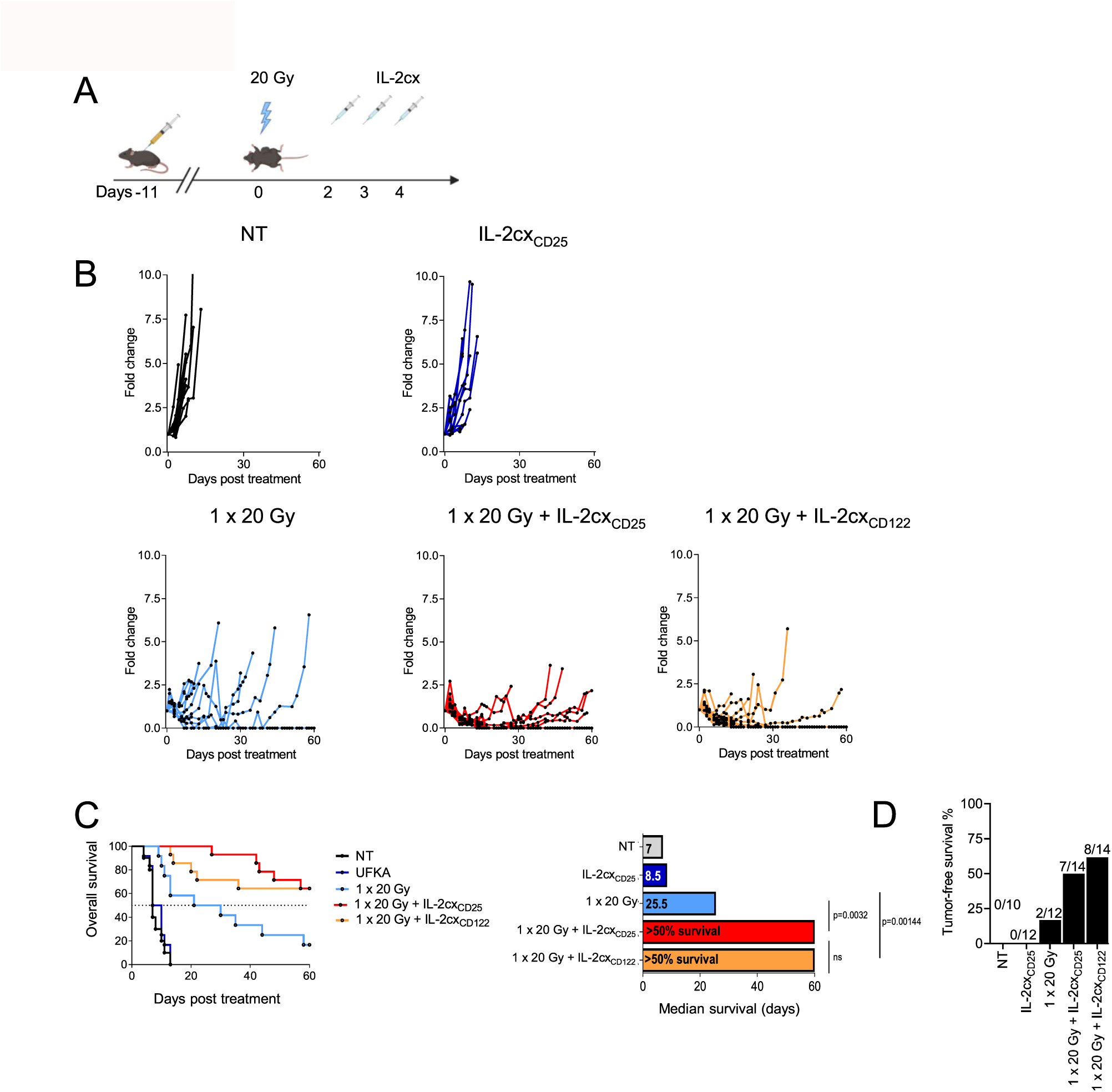
Combined radiotherapy and IL-2 immunotherapy efficiently controls primary tumors. **(A)** Schematic representation of the experimental set up. 10–11 days after tumor inoculation, mice were irradiated with 20 Gy, followed by treatment with IL-2 complexes (IL-2cx) 48 hours after irradiation for 3 consecutive days. **(B)** Tumor growth curves shown to day 60 post treatment. **(C)** Survival curve and median survival depicted. **(D)** Percentage of tumor free and tumor bearing mice at experiment termination. Data are presented as mean ± SEM of three to four independent experiments. Differences in median survival days were taken from the survival curves and analyzed by pairwise Log-rank (Mantel-Cox) test. IL-2cx_CD25_, CD25-biased IL-2/UFKAcx; IL-2cx_CD122_, CD122-biased IL-2/NARAcx; NT, non-treated.

To determine the impact of CD8^+^ T cells, we used a CD8-specific depleting mAb (**Fig. 3A**). Use of the CD8-specific depleting mAb abrogated the effects of both single-dose 20 Gy RT and IL-2cx_CD25_ and of single-dose 20 Gy RT and IL-2cx_CD122_ combination immunotherapies (**Fig. 3B**). These data indicated CD8^+^ T cells played a crucial role in combination therapy during the initiation of the anti-cancer immune response.

**Figure 3.**
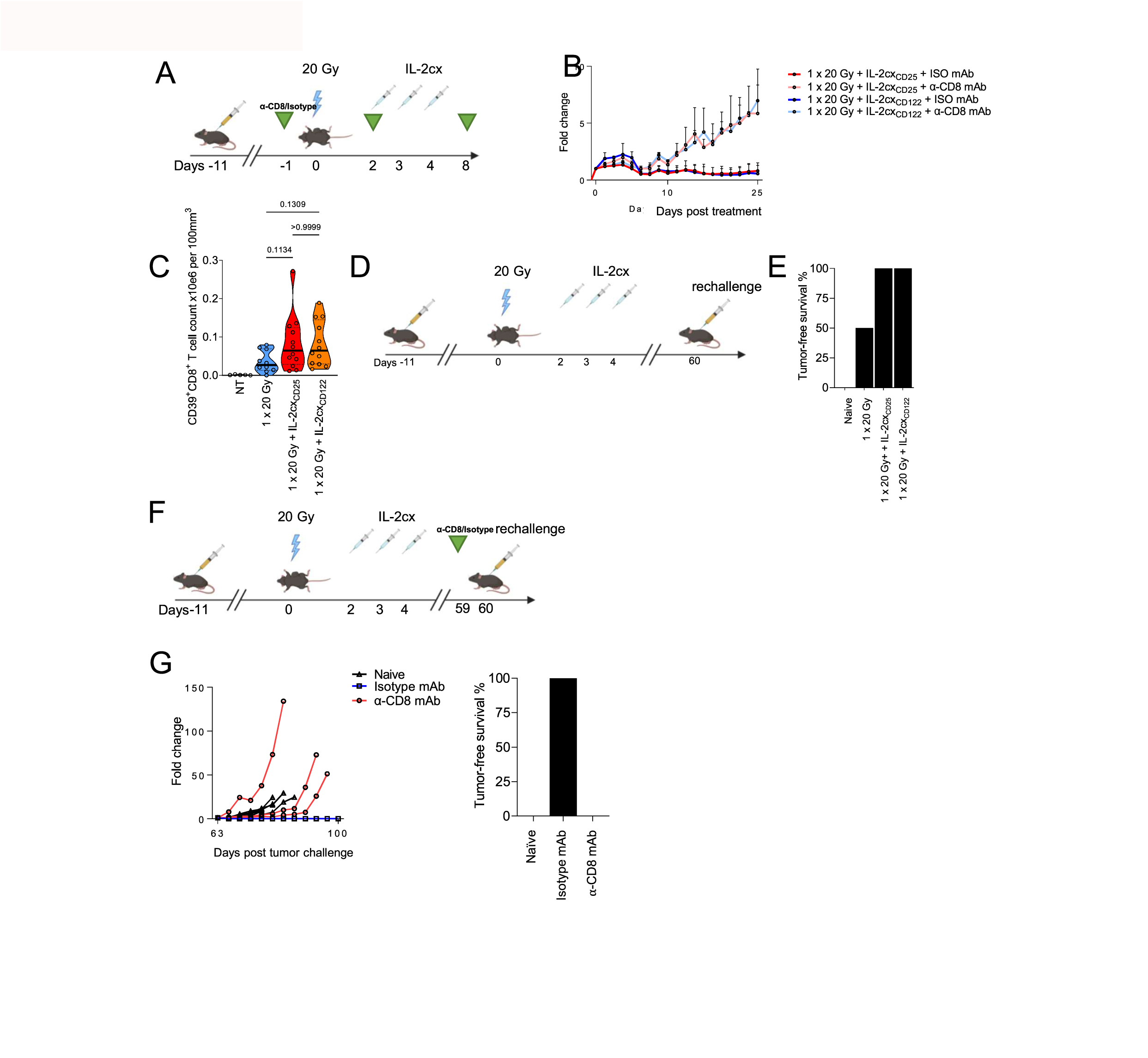
Combined radiotherapy and IL-2 immunotherapy generates long-lasting anti-tumor immune responses. **(A)** Mice were treated with a CD8-specific depleting monoclonal antibody (mAb) or an isotype-matched control mAb 24 hours prior to irradiation with two additional injections at 48 hours and 8 days after irradiation. **(B)** Tumor growth curves from experimental set up in A). **(C)** Quantification of intratumoral CD39^+^CD8^+^ T cells following treatment as in Fig. 2A. Tumors were harvested 9 days after irradiation and CD39^+^CD8^+^ T cells were determined by flow cytometry. **(D)** 10–11 days after tumor inoculation into the right flank, mice were irradiated with 20 Gy, followed by treatment with indicated IL-2cx 48 hours after irradiation for 3 consecutive days and monitoring until day 60. Mice with complete remissions were rechallenged with 5 x 10^5^ MC38 cells on the left flank and tumor growth was monitored until day 100 after initial treatment. Naïve non-tumor bearing mice were challenged as controls. **(E)** Percentage of tumor-free mice at experiment termination. **(F)** Tumor-bearing mice treated as in Fig. 2A and in complete remission received a CD8-specific depleting mAb or an isotype-matched control mAb on day 59 after treatment initiation and were rechallenged with with 5 x 10^5^ MC38 cells on the left flank. **(G)** Tumor growth in mice of (**F**), with percentage of tumor-free mice at experiment termination. Data are presented as mean ± SEM of one to three independent experiments. Differences in median survival days were taken from the survival curves and analyzed by pairwise Log-rank (Mantel-Cox) test.

Having observed a central role of tumor-infiltrating CD8^+^ T cells, we hypothesized that single-dose 20 Gy RT stimulated tumor antigen-specific CD8^+^ T cells. CD39 has been suggested to identify antigen-stimulated CD8^+^ T cells, rather than bystander-activated T cells, in tumors^36^. Quantification of intratumoral CD39^+^ CD8^+^ T cells showed that single-dose 20 Gy RT markedly increased counts of these cells in the tumor, which were further boosted by the addition of either IL-2cx_CD25_ or IL-2cx_CD122_ (**Fig. 3C**).

To assess the role of CD8^+^ T cells during the memory phase, mice that were in complete remission 60 days after RT were rechallenged with the same tumor cells on the opposing flank, followed by monitoring tumor growth (**Fig. 3D**). Whereas mice treated with RT combined with either IL-2cx were able to completely reject the rechallenging tumor cells, mice treated with RT alone or naïve mice that had never encountered tumor cells succumbed to the tumor rechallenge (**Fig. 3E**). When we administered a CD8-specific depleting mAb in the memory phase before rechallenging mice with tumor cells (**Fig. 3F**), mice lacking CD8^+^ T cells were unable to control the tumor rechallenge, whereas mice treated with an isotype-matched mAb remained tumor free until day 100 (**Fig. 3G**). Collectively, these data demonstrated that CD8^+^ T cells were essential both during initiation and long-term surveillance of the anti-cancer immune response following combined RT and IL-2cx immunotherapy.

### Combination radiotherapy and CD25-biased IL-2 immunotherapy favors expansion of tumor-infiltrating CD8^+^ T cells

We assessed whether the synergistic effect of RT and IL-2cx was localized to the irradiated tumors or also apparent systemically (**Fig. 4A**). Compared to untreated mice and animals receiving monotherapy or single-dose 20 Gy RT plus IL-2cx_CD122_, mice treated with single-dose 20 Gy RT in combination with IL-2cx_CD25_ harbored significantly higher percentages of intratumoral CD8^+^ T cells (**Fig. 4B**). Moreover, IL-2cx_CD25_ resulted in intratumoral (CD8/Treg) ratios that were significantly higher than those of the monotherapies or of untreated animals (**Fig. 4C**). These effects were also observed with IL-2cx_CD122_ when combined with RT, although they were less marked than with IL-2cx_CD25_. Contrary to what we observed in the tumors, IL-2cx_CD122_ combined with single-dose 20 Gy RT achieved the highest CD8/Treg ratios in DLNs, non-draining lymph nodes (NDLN), and spleen (**Fig. 4D**). Notably, the effects observed in these secondary lymphoid organs following treatment with IL-2cx_CD25_ alone or in combination with RT were very different and resulted in moderate changes or decreases of CD8/Treg ratios (**Fig. 4D**). These data suggested that IL-2cx_CD25_ activated CD8^+^ T cells preferentially in the tumor due to their CD25 upregulation following RT of the tumor, whereas IL-2cx_CD25_ stimulated CD8^+^ T cells in secondary lymphoid organs, i.e. extratumorally. This hypothesis was supported by significant expression of the proliferation marker Ki67 in tumor-infiltrating CD8^+^ T cells following RT and IL-2cx_CD25_, but not in DLNs, whereas the opposite was seen in CD8^+^ T cells of mice receiving RT plus IL-2cx_CD122_ (**Fig. 4E**).

**Figure 4.**
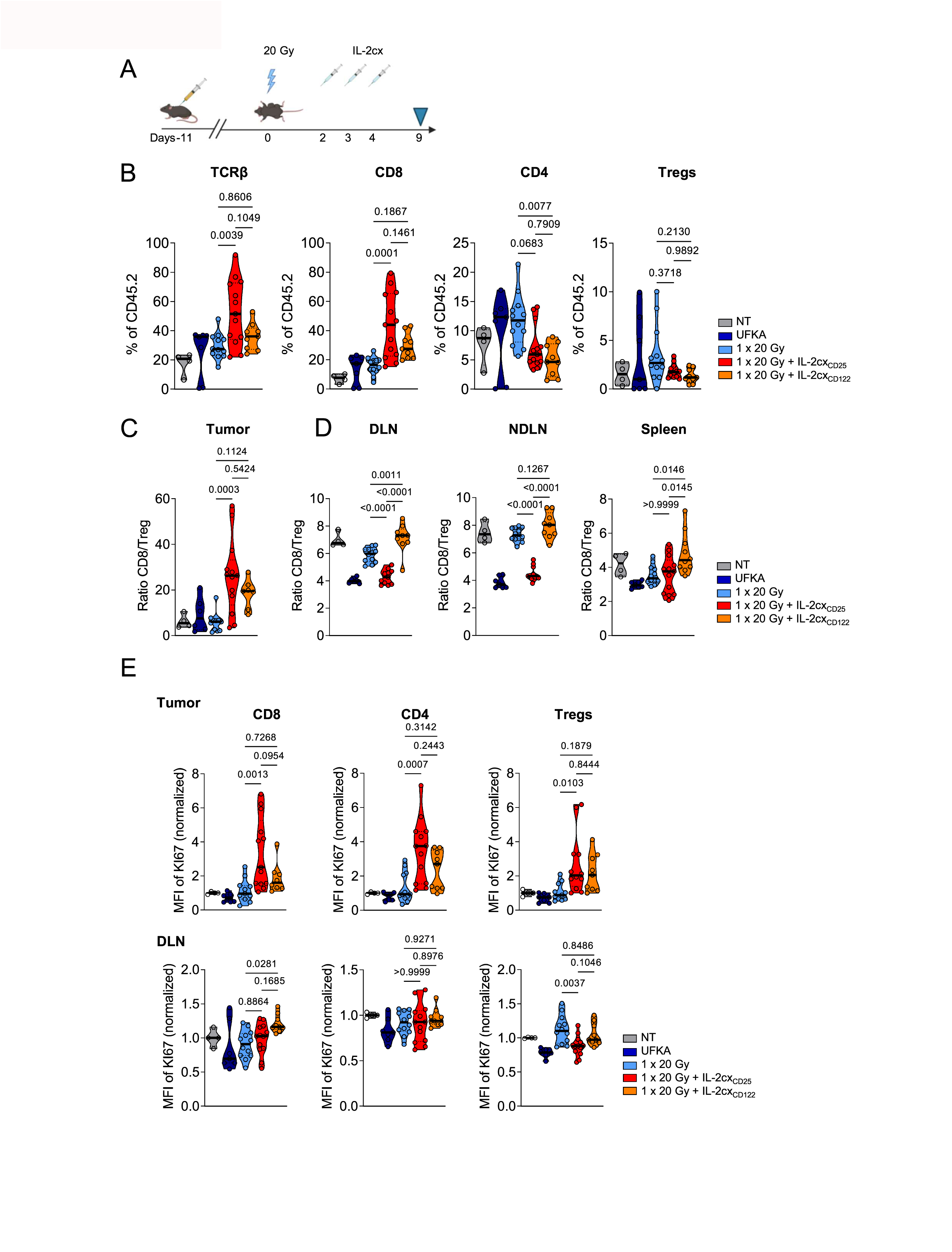
Combined radiotherapy and CD25-biased IL-2 immunotherapy expands tumor-infiltrating CD8^+^ T cells. **(A)** Schematic representation of the experimental set up. Experiment termination (harvest) denoted by blue triangle. **(B)** Percentages of TCRb^+^ T cells, CD8^+^ T cells, CD4^+^ T cells, and FoxP3^+^ Tregs in the tumor 9 days after irradiation. **(C)** Ratios of CD8^+^ T cells to Tregs in the tumor. **(D)** Ratios of CD8^+^ T cells to Tregs in the DLN, non-draining lymph node (NDLN) and spleen. **(E)** Mean fluorescence intensity (MFI) of Ki67 in CD8, CD4 and Tregs from tumor (top) and DLN (bottom). MFI normalised to the same immune subtype in non-treated mice. Data are presented as mean ± SEM of two to three independent experiments. Differences were analyzed using a one-way ANOVA.

### IL-2cx immunotherapy reverts radiotherapy-induced exhaustion in tumor-infiltrating CD8^+^ T cells

Given their crucial role, we hypothesized that IL-2cx could qualitatively affect intratumoral CD8^+^ T cells following combined RT and IL-2cx immunotherapy. As the immunosuppressive tumor microenvironment can induce T cell exhaustion, we assessed the transcription factor thymocyte selection-associated HMG BOX (TOX), which has been associated with exhausted T cells^37–39^, in tumor-infiltrating CD8^+^ T cells. Following RT, we observed a peak of both CD4^+^ and CD8^+^ T cell infiltration into the tumor at day 7 (**Fig. 5A**). This correlated with a rapid upregulation of TOX and PD-1 (**Fig. 5B and 5C**). Notably, combination of RT with either IL-2cx_CD25_ or IL-2cx_CD122_ immunotherapy reversed this exhaustion signature in tumor-infiltrating CD8^+^ T cells and significantly reduced TOX (**Fig. 5D**). These data demonstrated that IL-2cx treatment reinvigorated tumor-infiltrating CD8^+^ T cells following single-dose 20 Gy RT.

**Figure 5.**
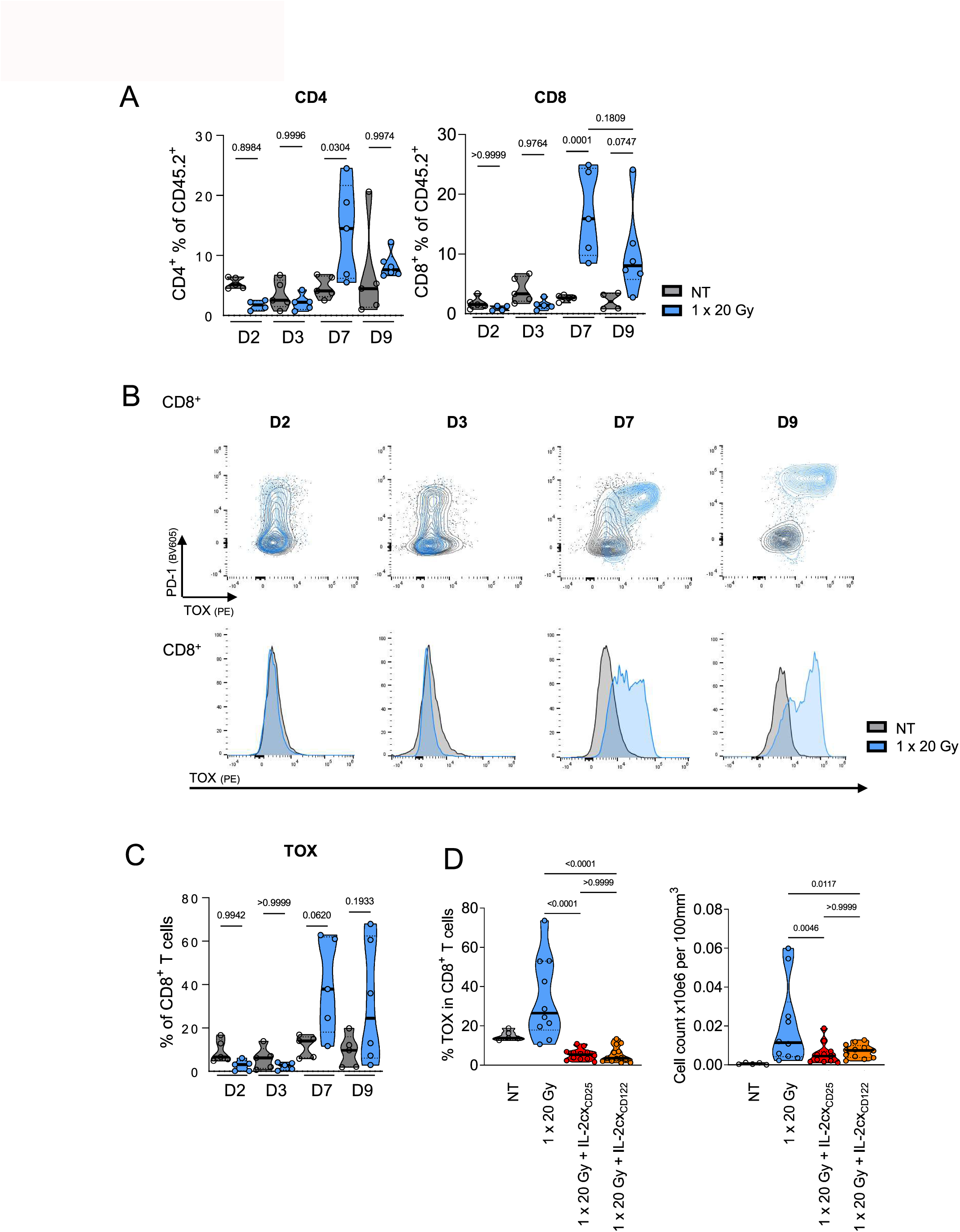
IL-2cx immunotherapy reverts radiotherapy-induced exhaustion in tumor-infiltrating CD8^+^ T cells. **(A)** Percentages of CD4^+^ and CD8^+^ T cells in CD45.2 tumor infiltrates at 2, 3, 7 and 9 days post irradiation with 20 Gy. **(B)** (Top) Representative FACs plots and (bottom) histograms gated on tumor infiltrating CD8^+^ T cells regarding PD-1 and TOX expression. **(C)** Percentage of TOX^+^ in CD8^+^ T cells over time. **(D)** Percentage and total number of TOX^+^ cells within (top) tumor-infiltrating CD8^+^ and (bottom) CD4^+^ T cells. Data are presented as mean ± SEM of three independent experiments. Differences were analyzed using a one-way ANOVA.

### Combined radiotherapy and IL-2 immunotherapy affects abscopal tumors

We observed slight differences in CD8^+^ T cells in tumors and secondary lymphoid organs following IL-2cx_CD25_ or IL-2cx_CD122_ immunotherapy combined with single-dose 20 Gy RT. Thus, we wondered whether these IL-2cx differed in their ability to induce systemic immune responses facilitating the control of distant, non-irradiated ‘abscopal’ tumors. To this end, we investigated the potency of our combined treatments in mice bearing two tumors (**Fig. 6A**). RT with 20 Gy alone was able to adequately control the primary irradiated tumor (**Fig. 6B**), however, it failed to have an impact on the non-irradiated abscopal tumor (**Fig. 6C**). The addition of either IL-2cx_CD25_ or IL-2cx_CD122_ not only improved control of primary tumors (**Fig. 6B**) but also of abscopal tumors (**Fig. 6C**). These effects resulted in prolonged overall survival of the animals, which increased from a median of 13 days in response to RT alone to 22 and 21 days when RT was combined with IL-2cx_CD25_ and IL-2cx_CD122_, respectively (**Fig. 6D**). This enhanced survival correlated with an increase in CD8/Treg ratios in abscopal tumors (**Suppl. Fig. 2**), similar to what we had observed for primary tumors (**Fig. 4C**). Collectively, combination of single-dose 20 Gy RT with IL-2cx immunotherapy induced potent anti-tumor immune reponses that acted on both primary and distant abscopal tumors.

**Figure 6.**
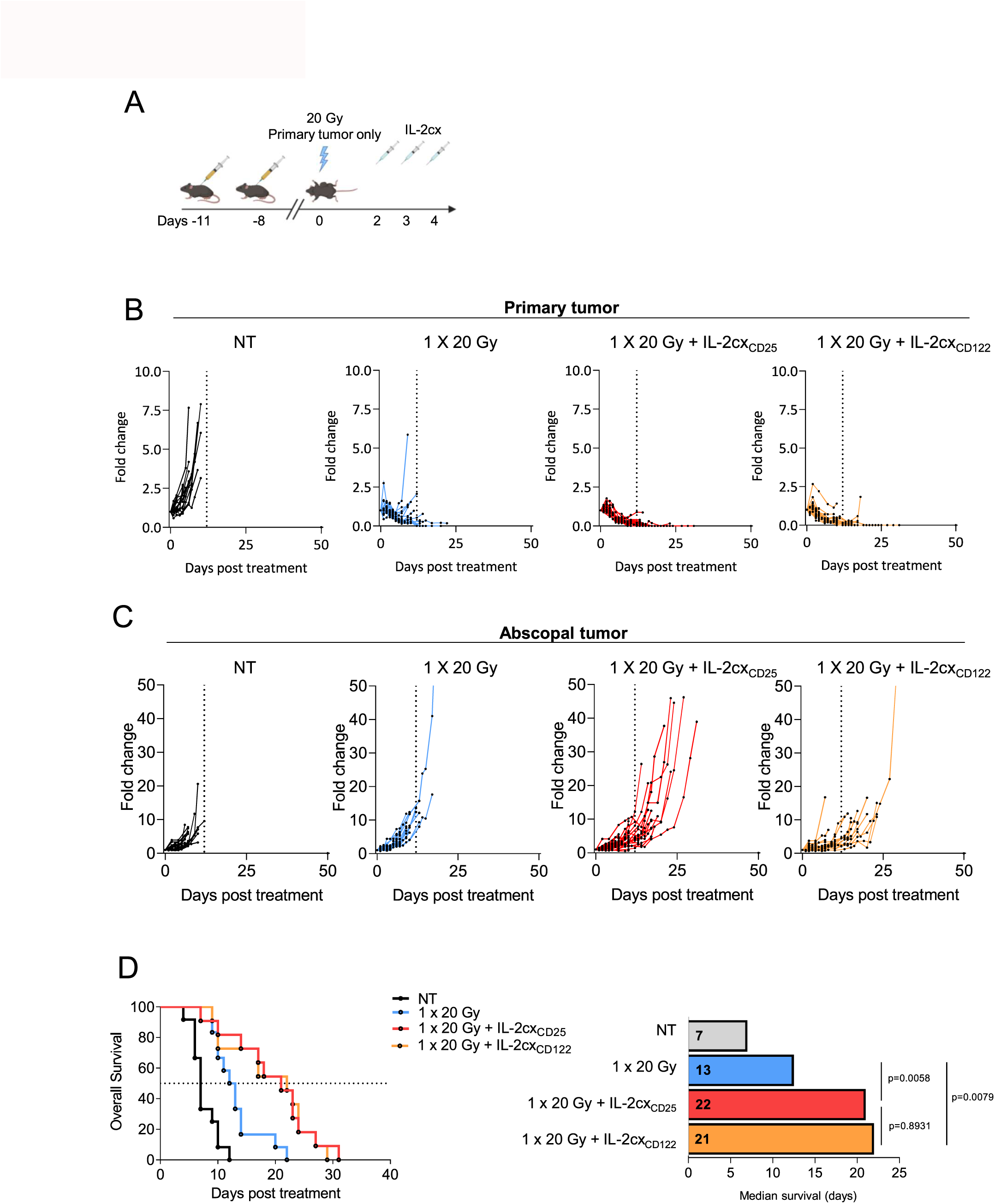
Combined radiotherapy and IL-2 immunotherapy affects abscopal tumors. **(A)** Schematic representation of the experimental set up. **(B)** Primary tumor growth curves. **(C)** Distant, non-irradiated tumor growth curves. **(D)** Survival of mice to a cumulative tumor burden of 1000 mm^3^ and median survival days post treatment. Data are presented as mean ± SEM of two to three independent experiments. Differences in median survival days were taken from the survival curves and analyzed by pairwise Log-rank (Mantel-Cox) test.

## Discussion

In this study, we found that single-dose 20 Gy RT increased CD25 and, to a lesser extent, also CD122 on tumor-infiltrating CD8^+^ T cells, which allowed us to combine this RT scheme with a CD25-biased (made with UFKA-20 mAb) and a CD122-biased IL-2cx (using NARA1 mAb). Such single high-dose RT plus IL-2cx combination immunotherapy relied on long-lived CD8^+^ T cells, resulted in efficient anti-tumor immune responses against both primary and abscopal tumor nodules, and formed long-lasting immune memory. Interestingly, RT-induced exhaustion of tumor-infiltrating CD8^+^ T cells was reversed by use of either IL-2cx. Several aspects of our findings are interesting and novel when compared to previously published data, as discussed below.

The effect of RT and the subsequent immune changes that ensue can have a major impact on not only the type of immunotherapy that can be applied but more importantly the overall outcome of the response. Differential fractionation regimens and dosages that theoretically may induce similar levels of toxic DNA damage, can however result in very different immunological outcomes, and we are still lacking a rational approach to optimally combine RT with molecularly-designed immunotherapeutic agents. Furthermore, timing and sequencing of additional immunotherapies is essential, given the targets of immunotherapy are generally expressed in a kinetic manner after RT^40^. Whereas the current gold standard in immune response-inducing RT is the use of a hypofractionated scheme^5,41^, which is derived from a small number of empirical preclinical studies, we and others have observed favorable therapeutic responses when using a single high dose of 20 Gy of RT in conjunction with immunotherapeutic agents^35^. These differences may result from the heterogeneity of tumor models and types of immunotherapy that are used^6^, however, further head-to-head analysis comparing different dosages and fractionations in combination with immunotherapies is warranted^40^.

We observed that CD25 expression on tumor-infiltrating CD8^+^ T cells peaked two days after single-dose 20 Gy RT. This finding and kinetic is similar to CD25 upregulation on antigen-specific CD8^+^ T cells following viral infection of mice^42^. This increase in CD25 expression on tumor-infiltrating CD8^+^ T cells shortly after RT allowed us to strategically target recently-activated CD8^+^ T cells using a CD25-biased IL-2cx. The expansion of polyclonal T cells through irradiation allows for a much more relevant clinical setting, whereby immunologically ‘cold’ tumors can be turned ‘hot’ through RT. The utilization of consequential immunotherapies is therefore reliant on the release of neo-antigens and the subsequent *de novo* priming of immune responses^43,44^. Previous studies using CD25-biased IL-2 formulations have relied on TCR-restricted T cells^45,46^, which often have a much higher affinity for their antigen and thus generate a much stronger anti-tumor response. These are often adoptively transferred with concomitant adjuvants, such as lipopolysaccharide, in addition to IL-2 formulations to maximize the anti-tumor response. As such, our model system more closely mimics a translational setting, whereby polyclonal T cells are activated to upregulate CD25 in the absence of further adjuvant stimulation and relies on the priming with endogenous tumor antigens to generate an efficient immune response.

Despite the fact the levels of CD25 present on recently-activated CD8^+^ T cells in the tumor were almost 10-fold less than that of Tregs, we observed an impressive expansion of recently-activated intratumoral CD8^+^ T cells by about 30–40 fold upon RT plus IL-2cx immunotherapy compared to about 16 fold in RT alone, with only minimal Treg expansion with both IL-2cx_CD25_ and IL-2cx_CD122_. Strikingly, the CD8/Treg ratios within the tumor were roughly 25 fold, which was comparably larger than that of another pre-clinical study using IL-2cx ^45^. Other publications using CD122-biased IL-2 formulations have reported enhanced CD8^+^ T cell proliferation and CD8/Treg ratios in secondary lymphoid organs^20,23,30,33^, which can result from the large numbers of memory CD8^+^ T cells present in secondary lymphoid organs and responding to CD122-directed IL-2 formulations. Such extratumoral CD8^+^ T cell expansion was recently also demonstrated in multiple tumor models upon treatment with a differential IL-2cx_CD122_ combined with RT^30^ and correspond to our own results with RT in combination with CD122-biased IL-2/NARA1cx, but not with CD25-biased IL-2/UFKAcx. Of note, the epitope-directed approach to increase the concentration of IL-2 in the tumor as part of a fusion protein (L19-IL2), which was successfully combined with single high-dose RT, also resulted in intratumoral CD8^+^ T cell expansion with only minimal changes in secondary lymphoid organs^47^. However, its application is restricted to tumors with neovasculature expressing the ED-B-domain epitope. As we detected CD25 upregulation only in tumor-infiltrating CD8^+^ T cells after single-dose 20 Gy RT, we observed a strong synergistic effect with IL-2cx_CD25_ in the irradiated tumor, with minimal changes observed in peripheral organs. This correlated with the highest Ki67 abundance in intratumoral CD8^+^ and conventional CD4^+^ T cells, whereas proliferation of these cells following single-dose 20 Gy RT and IL-2cx_CD25_ was negligible in secondary lymphoid organs. Conversely, a combination treatment of single-dose 20 Gy RT and IL-2cx_CD122_ resulted in expansion of CD8^+^ T cells both in the tumor and in secondary lymphoid organs.

The presence of CD8^+^ T cells in the therapeutic response to radioimmunotherapy is well established to be an integral component. Similar to other studies^48^, we demonstrated the necessity for CD8^+^ T cells in this anti-tumor response. Depletion of CD8^+^ T cells prior to tumor inoculation or prior to rechallenge during the memory phase completely ablated tumor control. Thus, we observed no anti-tumor response in the absence of CD8^+^ T cells in both RT plus IL-2cx_CD25_ and RT plus IL-2cx_CD122_ treatment groups. This indicated that CD25 and CD122 expression on non-CD8^+^ T cells, including innate lymphoid cells and NK cells^15,22^, was negligible in this tumor model.

Modified IL-2 variants such as fusion proteins or antibody conjugates targeting CD122 have already demonstrated anti-tumor responses in preclinical models^20,22–24,30^. In addition to stimulating expansion, IL-2 is also an integral factor in memory programming during the initial priming phase of T cells^49,50^. In our hands, leaving out IL-2cx during treatment failed to generate a long-term T cell memory, indicating the requirement for IL-2 signals during the priming phase of the immune response^51^. Indeed, in the absence of IL-2cx, RT alone upregulated markers correlating with T cell exhaustion, such as TOX and PD-1. While TOX, when co-expressed with other co-inhibitory markers, has been associated with an exhausted phenotype as a result of chronic antigen stimulation, such as in viral infections or cancer^37,38^, TOX expression is also regulated by the inflammatory environment^52^. Similar to previous findings^23,46^, we also observed that co-administration of IL-2cx induced a reduction in PD-1 expression, rendering cells less vulnerable to immunosuppression by this pathway and therefore sustained for longer periods in the tumor. Our study focused mainly on the two markers of PD-1 and TOX to define an exhausted phenotype in tumor-infiltrating CD8^+^ T cells. Indeed the classification of ‘exhaustion’ includes a much broader scope of markers (reviewed in ^53,54^), however the transcription factor TOX plays in important role in defining T cell exhaustion ^37,38^. A broader panel of transcription factors and co-inhibitory molecules, including T cell factor 1 (TCF-1), could have provided a more granular view of the different stages of exhaustion of tumor-infiltrating CD8^+^ T cells ^55,56^. The use of IL-2 variants has also been recently shown to rescue cells from terminal differentiation^31,57^. IL-2-induced reduction of exhaustion markers may therefore be due to its role in preventing tumor-infiltrating CD8^+^ T cells from commitment to a terminally differentiated fate. Indeed two recent publications studying chronic viral infection and showed the use of IL-2 immunotherapy in combination with PD-1 antagonism synergized to alter the differentiation program of precursors of exhausted CD8^+^ T cells – also termed stem-like CD8^+^ T cells characterized by co-expression of PD-1 and TCF-1 – to develop into a distinct and highly functional effector T cell population^58,59^. Alternatively, by inducing a larger expansion of effector CD8^+^ T cells may allow for a faster eradication of the tumor and therefore reducing the amount of available antigen stimuli present. Further experiments will be required to gather in-depth knowledge of the role of IL-2 in resetting irradiation-induced TOX and TCF-1 expression in intratumoral CD8^+^ T cells.

The combination of RT and modified IL-2 variants has achieved potent anti-cancer responses in both preclinical models as well as in clinical trials and is proving to be an efficacious therapeutic combination^32,33,60^. However, a majority of these IL-2 variants target the dimeric IL-2R to stimulate memory CD8^+^ T cells and avoid Treg expansion. Our data demonstrate for the first time to our knowledge the combined use of RT with an IL-2cx_CD25_. Although both IL-2cx_CD25_ and IL-2cx_CD122_ achieved similar anti-tumor responses in our preclinical model, our data suggest that IL-2cx_CD25_ preferentially stimulated recently-activated intratumoral CD8^+^ T cells, whereas IL-2cx_CD122_ expanded antigen-experienced memory CD8^+^ T cells in both secondary lymphoid organs and the tumor microenvironment.

## Methods

### Cell line

The MC38 murine colorectal cancer cell line was a gift by Lubor Borsig, Department of Physiology, University of Zurich, Switzerland. Cells were cultured in RPMI (Gibco) supplemented with 10% fetal bovine serum (FBS, Gibco), 1% penicillin/streptomycin (Gibco) and 1% Glutamax (Gibco) at 37°C and 5% CO2. A fresh aliquot of frozen cells was used per experiment. Cell lines were passaged at least 3 times prior to injection into mice and were kept no more than 2 weeks in culture.

### Mice

All animal experiments were performed according to Swiss federal and cantonal law and approved by the Cantonal Veterinary Office Zurich (ZH113/2020). 8–10-week old female C57BL/6J mice were purchased from Envigo. Mice were allowed to acclimatize at least one week prior to start of experiment.

### Tumor model

Cells were thawed and passaged at least 3 times before injection. Mice were anesthetized with isoflurane (Attane, Piramal Ltd.) at a rate of 1 L/min oxygen with 3.5% isoflurane during inoculation. Mice were shaved on the flank in which the tumor was to be inoculated. 5–7.5 x 10^5^ cells were resuspended in 100 μl of phosphate buffered saline (PBS) and injected subcutaneously on the right flank of the mouse. Abscopal tumors were inoculated in the same fashion 3 days later on the left flank of the mouse. Treatment started 10–11 days after inoculation of the primary tumor. Tumors were measured with calipers and mice were randomized into treatment groups on the day according to the tumor volume (both primary and abscopal). Tumor volume was calculated as follows: V=L*W^2^/2, where V is volume, L is length and W is width. For tumor rechallenge experiments, cured mice were inoculated with 5 x 10^5^ MC38 cells on the opposing flank of the original primary tumor. Survival was defined as tumor growth to 500 mm^3^ for the primary tumor model or a cumulative 1000 mm^3^ for the bilateral tumor model.

### Irradiation

RT treatment was performed as previously described^61^. In brief, we used the X-RAD SmART (Precision X-Ray Inc.) small animal image-guided radiation research platform. Mice were anesthetized during the entire procedure. Tumor irradiation was performed with a 8 x 12 mm beam with the isocenter placed in the middle of the tumor. Target planning of irradiation was performed to ensure minimal exposure to all areas surrounding the tumor. As indicated, a dose of 8 Gy or 20 Gy was delivered to the tumor with care to avoid irradiation of the inguinal (draining) lymph node. Treatment planning was performed using a dedicated small animal treatment planning software SmART-ATP (SmART Scientific Solutions B.V.).

### IL-2 complexes

Recombinant human IL-2 (Proleukin) was purchased from R&D Systems or obtained from the National Cancer Institute’s Biological Research Branch. IL-2 complexes (IL-2cx) were generated, as previously described^18,20^, by mixing 1 μg IL-2 with 10 μg monoclonal antibody (mAb) at a 1:1 molar ratio, resulting in CD25-biased IL-2/UFKA-20 complexes (IL-2cx_CD25_) or CD122-biased IL-2/NARA1 complexes (IL-2cx_CD122_). IL-2cx were delivered in PBS via intraperitoneal injection 48 hours after irradiation for three consecutive days.

### *In vivo* depletion

Where indicated, 300 μg of depleting anti-mouse CD8 mAb (clone YTS169, BioXcell) or an IgG2a isotype control mAb (C1.18.4, BioXcell) were resuspended in 200 μL PBS and injected intraperitonally one day prior to irradiation as well as two and eight days after irradiation, or one day prior to tumor rechallenge (day 59). Depletion efficiency was confirmed 24 hours after injection by flow cytometry on tail vein blood.

### Flow cytometry

Tumors, spleens and lymph nodes were harvested and processed, as published^20,22,23^. In brief, tumors were cut into pieces with scissors and incubated on a rolling platform in 3 ml/sample of tumor digestion buffer consisting of complete tissue culture media plus 200 U/ml collagenase D for 45 minutes at 37°C. Red blood cells in blood and spleen samples were removed using ammonium-chloride-potassium lysis buffer. All organs were mechanically dissociated, processed into single cell suspensions using a 70-μm cell strainer and stained with an extracellular staining mix, as previously established^62^. Subsequently, cells were fixed and permeabilized with the Foxp3/Transcription Factor Staining Buffer Set (eBioscience) according to the manufacturer’s instructions. Intracellular staining was performed overnight or for a minimum of an hour. **Supplementary Table 1** provides a list of all antibodies used for flow cytometry. Samples were acquired on a Fortessa (DIVA) or Cytek Aurora (Cytek) flow cytometer and analyzed with FlowJo software v10.8 (BD Biosciences).

### Statistical analysis

Mice were randomized based on tumor volume and assigned to experimental groups. Data are displayed as mean ± standard error of the mean. Statistical analysis was performed in GraphPad Prism v9.3. Outliers were assessed using the outlier test in Prism and excluded accordingly. Comparisons of flow cytometry data were performed using a one-way ANOVA. Comparisons of survival curves were performed using Log-rank (Mantel-Cox) test. Statistical significance was established at *P* < 0.05. Figures were generated with BioRender.

## Acknowledgments

We thank Laura Bürgi, Dilara Sahin, Rebecca Masek and Lucas Schmid for help with experiments and the Pruschy and Boyman laboratories for helpful discussions.

## Author contributions

C.S.M.Y. designed and performed the experiments, analyzed the data, and wrote the manuscript. I.T. and L.G. performed experiments. M.E.R. provided essential data. M.P. and O.B. conceived the project, designed and analyzed experiments, and wrote the manuscript. All authors edited and approved the final manuscript.

