## Supplementary Figures and Table for "IL-2 immunotherapy rescues irradiation-induced T cell exhaustion *in vivo*"

**Supplementary Figure 1. Irradiation reduces CD25 and Foxp3 expression in tumor-infiltrating regulatory T cells.** (Top) Representative FACS plots gated on tumor-infiltrating CD4$^{+}$Foxp3$^{+}$ regulatory T cells (Tregs) at day 2 post irradiation. (Bottom) Percentage of Tregs in CD45.2 infiltrates and MFI of CD25 and Foxp3 expression normalized to non-treated mice. Data are presented as mean ± SEM of two to three independent experiments. Differences were analyzed using a one-way ANOVA.

**Supplementary Figure 2. A favorable CD8/Treg ratio is observed in non-irradiated abscopal tumors after combination therapy.** Ratios of CD8$^{+}$ T cells to Tregs in non-irradiated abscopal tumors. Data are presented as mean ± SEM of three independent experiments. Differences were analyzed using a one-way ANOVA. IL-2cx$_{CD25}$, CD25-biased IL-2/UFKAcx; IL-2cx$_{CD122}$, CD122-biased IL-2/NARAcx; NT, non-treated.

**Supplementary Table 1. Antibodies used for flow cytometry.**

| **Antigen** | **Fluorochrome** | **Clone** | **Manufacturer** | **Dilution** |
|---|---|---|---|---|
| CD122 | APC | TM-β1 | BioLegend | 1:300 |
| CD25 | PE | PC61 | BioLegend | 1:300 |
| CD39 | PerCP-eFluor710 | 24DMS1 | Invitrogen | 1:300 |
| CD4 | BUV496 | GK1.5 | BD | 1:400 |
| CD44 | BV510 | IM7 | BD | 1:400 |
| CD45.2 | Alexa Fluor 700 | 104 | BioLegend | 1:250 |
| CD8 | APC | 53-6.7 | BioLegend | 1:400 |
| Fixable viability dye | eFluor780 | | eBioscience | 1:1000 |
| Foxp3 | PE-Cy7 | FJK-16s | Invitrogen | 1:200 |
| Ki67 | BV605 | 16A8 | BioLegend | 1:200 |
| NK1.1 | BV711 | PK136 | BioLegend | 1:400 |
| PD-1 | BV605 | 29F.1A12 | BioLegend | 1:300 |
| TCRβ | BUV563 | H57-597 | BD | 1:400 |
| TOX | PE | TXRX10 | Invitrogen | 1:200 |

### Supp. Fig. 1

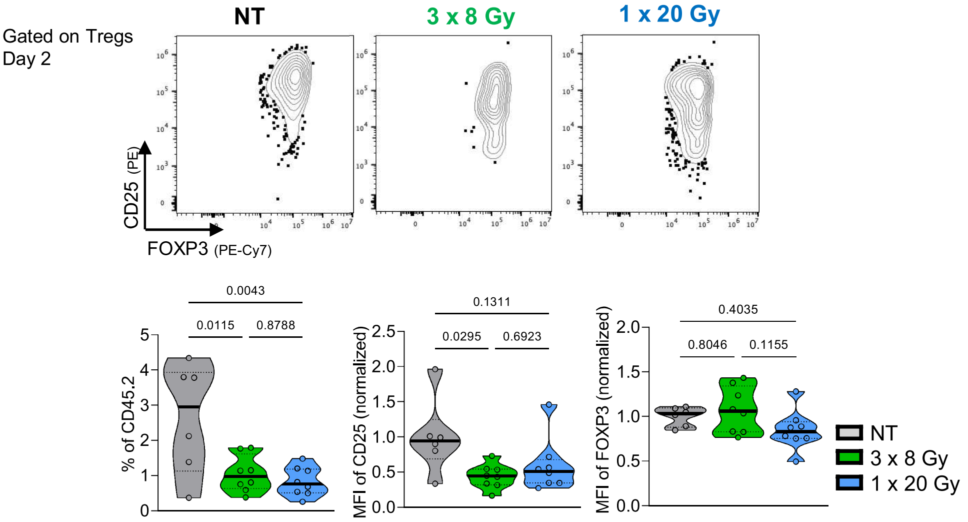


### Supp. Fig. 2

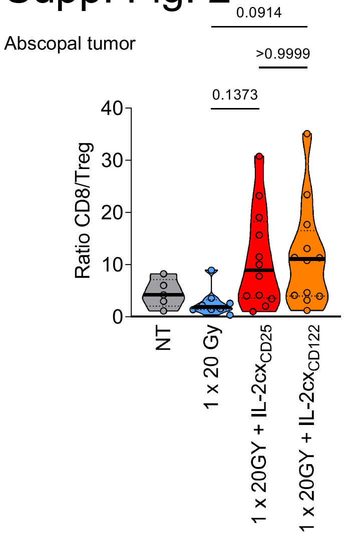
